# Mesothelial cells are not a source of adipocytes in mice

**DOI:** 10.1101/2021.05.13.444022

**Authors:** Gregory P. Westcott, Margo P. Emont, Jin Li, Christopher Jacobs, Linus Tsai, Evan D. Rosen

## Abstract

Visceral adipose tissue (VAT) depots are associated with the adverse metabolic consequences of obesity, such as insulin resistance. The developmental origin of VAT depots and the identity and regulation of adipocyte progenitor cells have been active areas of investigation. In recent years, a paradigm of mesothelial cells as a source of VAT adipocyte progenitor cells has emerged based on lineage-tracing studies using the Wilms’ tumor gene, *Wt1*, as a marker for cells of mesothelial origin. Here we show that *Wt1* expression in adipose tissue is not limited to the mesothelium, but is also expressed by a distinct preadipocyte population in both mice and humans. We identify keratin 19 (*Krt19*) as a highly-specific marker for the adult mouse mesothelium, and demonstrate that *Krt19*-expressing mesothelial cells do not differentiate into visceral adipocytes. These results contradict the assertion that the VAT mesothelium can serve as a source of adipocytes.

## Introduction

Adipogenic precursor cell populations have been proposed to arise from the mural cell compartment of the adipose vasculature(Tang et al., 2008), the endothelium (Tran et al., 2012), the bone marrow (Rydén et al., 2015) and from a subset of Sca-1+CD24+ (Rodeheffer et al., 2008) or Pdgfra+ (Rodeheffer et al., 2008) stromal cells. In addition, a mesothelial source of visceral adipocytes was hypothesized based on the established function of WT1 in mesenchymal development (Hastie, 2017), its presence in VAT and its absence in other white adipose depots (Chau et al., 2011), and its importance in the development of tissues such as the epicardium (Zhou et al., 2008). Chau et al. used fate mapping techniques to demonstrate that *Wt1*-expressing cells differentiate into mature adipocytes in murine VAT (Chau et al., 2014). Importantly, the conclusion that traced adipocytes are of mesothelial origin depends on exclusive expression of *Wt1* in mesothelial cells. In this study, we demonstrate that *Wt1* is expressed in a distinct preadipocyte population in addition to the mesothelium, and that mesothelial cells tracked with a more specific marker do not undergo adipocyte differentiation.

## Results

### Wt1 is not specific to visceral adipose mesothelium, and is also expressed in a distinct preadipocyte population

The advent of single-cell RNA-sequencing (scRNA-seq) technology has enabled the rigorous interrogation of previously accepted cell-type specific markers as well as the discovery of new ones. We performed scRNA-seq on adult male murine epididymal adipose tissue depots (**Fig. 1a**) and identified distinct mesothelial and preadipocyte populations (**Fig. 1b**). While established adipocyte progenitor markers such as *Pdgfra* and Sca-1 are present exclusively in the preadipocyte cluster, and mesothelial markers *Msln* and *Upk3b* are present exclusively in the mesothelial cluster, *Wt1* is strongly expressed in both populations (**Fig. 1c-d**). Expression of *Wt1* in both mesothelial and preadipocyte clusters was also noted in human omental single-nucleus RNA-sequencing data (**Fig. S1**). In contrast, our data indicate that *Krt19* is a highly specific marker for the mesothelium in both mouse and human **(Fig. 1c-d, S1)**. A member of the keratin family, KRT19 is an intermediate filament protein discovered in squamous cell carcinomas and found in epithelioid cells (Wu and Rheinwald, 1981). *KRT19* is expressed in human omental mesothelium but is absent from subcutaneous fat (Darimont et al., 2008).

**Figure 1.**
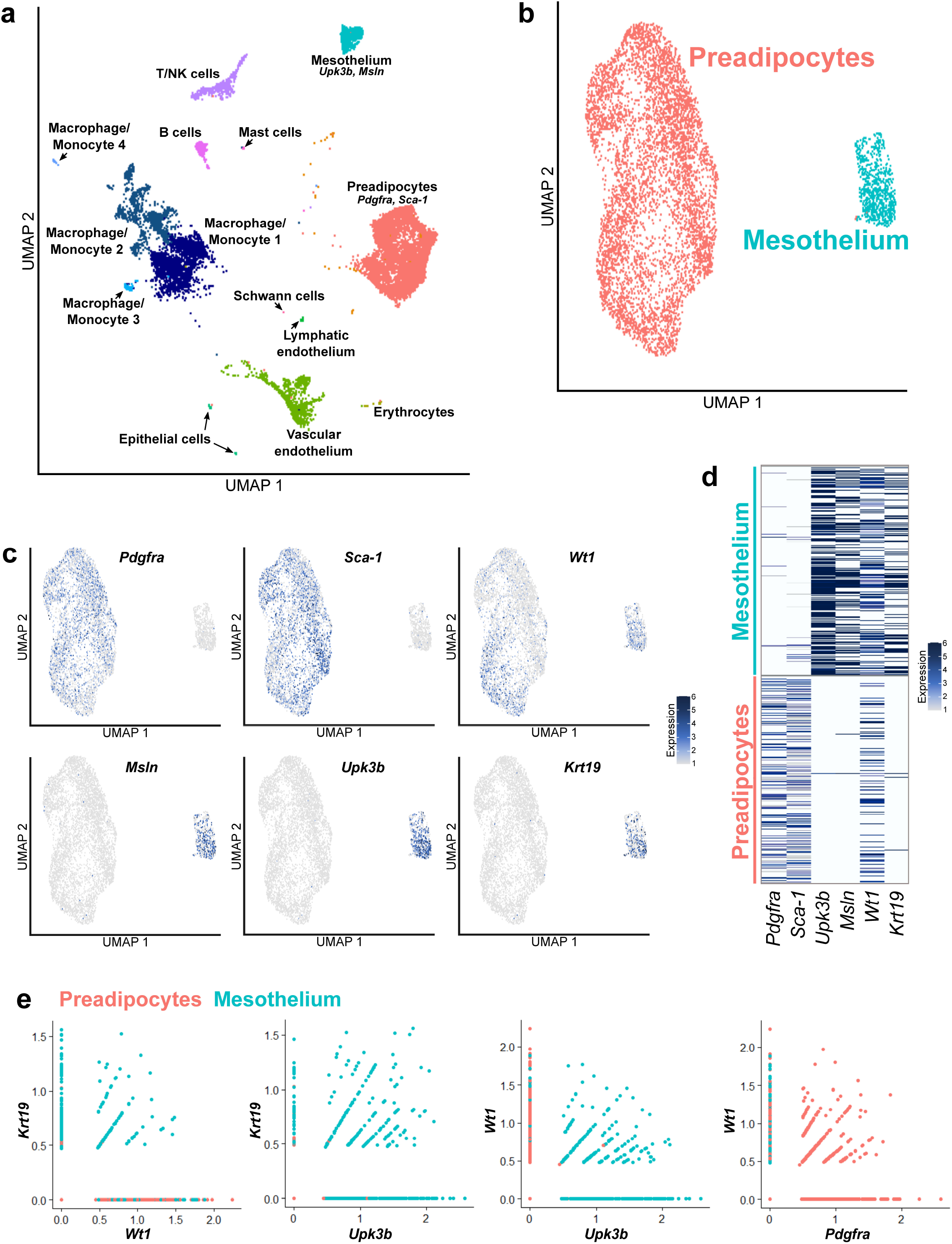
Single cell RNA-seq of mouse SVF reveals distinct preadipocyte and mesothelial populations which both express Wt1. **a**, UMAP plot of single cell RNA-seq performed on the SVF of mouse eWAT, which identifies all expected populations, including distinct populations of preadipocytes and mesothelial cells. **b**, UMAP plot of the preadipocyte and mesothelial populations. **c**, The preadipocyte population is characterized by the expression of *Pdgfra* and *Sca-1*, while the mesothelium expresses *Upk3b* and *Msln*. While *Krt19* expression mirrors that of mesothelial markers, *Wt1* is expressed in both preadipocytes and mesothelial cells. **d**, Heat map demonstrates that *Wt1* is expressed in cells expressing both preadipocyte and mesothelial genes. **e**, Most cells expressing *Wt1* but not *Krt19* are preadipocytes, while virtually all *Krt19*-expressing cells are mesothelial. A significant subset of *Wt1*+ cells co-express *Pdgfra*.

Trypsin digestion (Kenny et al., 2007) of intact fat pads was used to enzymatically isolate the mesothelium from perigonadal VAT depots, and qPCR for the mesothelial genes *Msln, Upk3b*, and *Krt19* confirmed high enrichment in the mesothelial fraction (**Fig. 2a**). *Wt1*, however, was expressed in both the mesothelium and the residual stromal vascular fraction (SVF). In order to isolate *Wt1*-expressing and *Krt19*-expressing cells, we obtained tamoxifen-inducible Cre mouse models for *Wt1* (Wt1^CreERT2^) (Zhou et al., 2008) and *Krt19* (Krt19^CreERT^) (Means et al., 2008), and crossed them to a reporter strain which expresses tdTomato following Cre recombination. We next performed flow cytometric analysis of Wt1^CreERT2^: and Krt19^CreERT^:tdTomato+ SVF cells stained for PDGFRA, and demonstrated that Wt1^CreERT2^: tdTomato+ cells display extensive co-labelling for PDGFRA while Krt19^CreERT^:tdTomato+ cells did not (**Fig. 2b**). Likewise, CD45-PDGFRA+ cells sorted from wild type SVF express *Wt1*, but not the other, more specific, mesothelial markers (**Fig. 2c**). Using our Wt1^CreERT2^: and Krt19^CreERT^:tdTomato+ models, we confirmed that trypsin-digested mesothelium is labeled by both models, but only Wt1^CreERT2^ also labels a substantial proportion of the remaining SVF population (**Fig 2d**).

**Figure 2.**
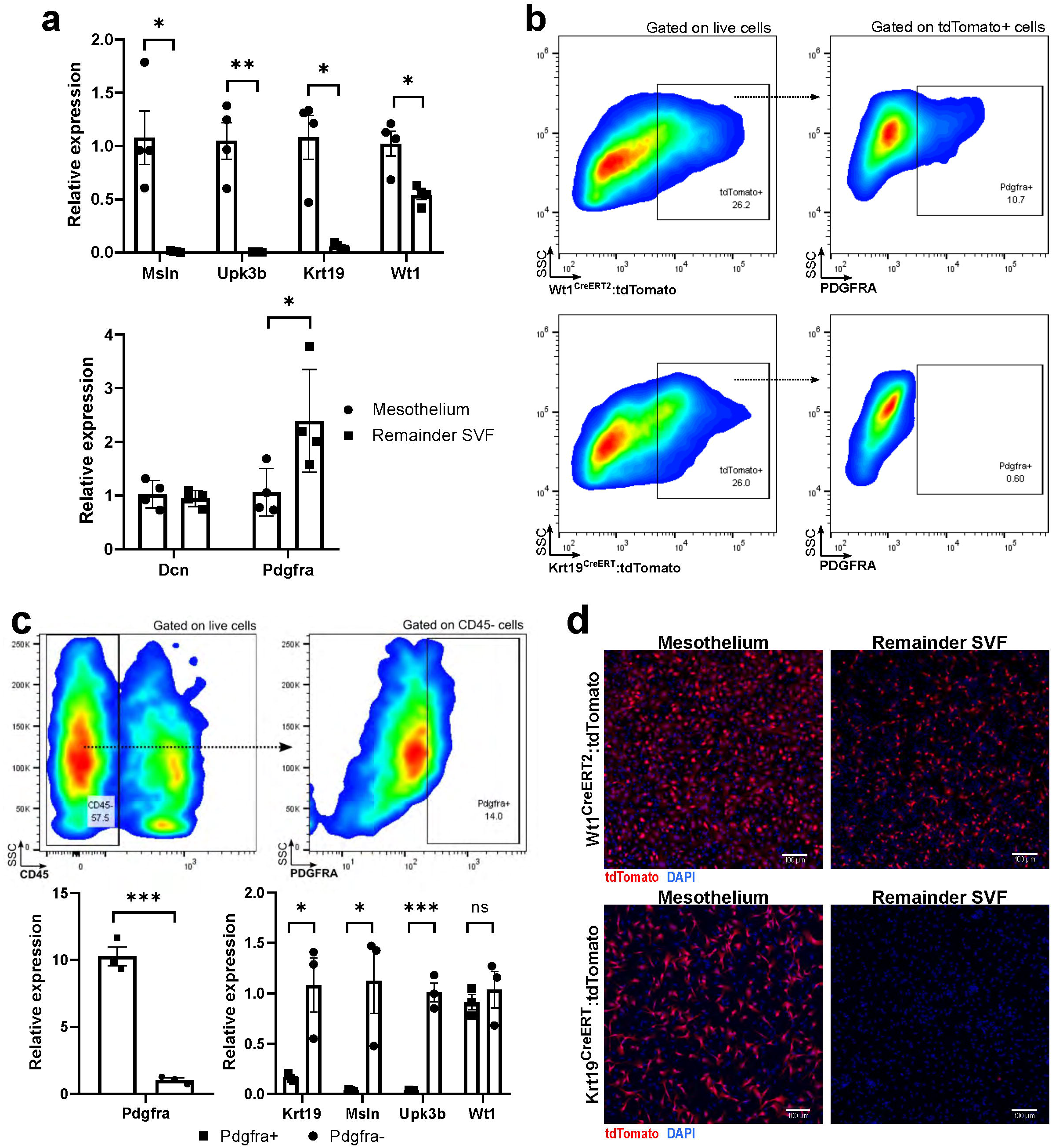
Wt1 is not a specific marker for the mesothelium and also labels Pdgfra+ preadipocytes. **a**, The mesothelium can be digested away from the remainder of the adipose parenchyma with trypsin, a technique that highly enriches for mesothelial markers, but not for Wt1, which is also highly expressed in the remainder SVF. The remainder SVF highly expresses *Pdgfra*, while *Dcn* is expressed by both mesothelium and other stromal cells. **b**, Using Wt1^CreERT2^ and Krt19^CreERT^ models crossed with a tdTomato reporter, flow cytometry reveals that *Wt1*-expressing cells contain a PDGFRA+ preadipocyte population, while *Krt19*-expressing cells do not. **c**, SVF cells isolated from wild-type mice were gated against CD45 and sorted into PDGFRA+ and PDGFRA-populations; mesothelial gene expression was restricted to PDGFRA-populations, while *Wt1* was also present in the PDGFRA+ preadipocyte population. **d**, Trypsin-digested mesothelium contains a high proportion of tdTomato+ cells in both Krt19^CreERT^ and Wt1^CreERT2^ models, while the remainder SVF contains tdTomato+ cells in only the Wt1^CreERT2^ mice. Error bars represent the standard error of the mean. ns, p ≥ 0.05, *, p <0.05; **, p<0.01; ***, p <0.001. SSC: side scatter.

### Cultured mesothelial cells do not undergo adipocyte differentiation *in vitro*

We next assessed the ability of Wt1+ and Krt19+ cells to undergo adipogenesis *ex vivo*. Wt1^CreERT2^: and Krt19^CreERT^:tdTomato+ mice were injected with tamoxifen and a week later VAT was harvested, collagenase-digested, and the SVF cultured in adipocyte differentiation media. While many newly formed adipocytes were tdTomato+ when Wt1^CreERT2^ was used as the driver, virtually no tdTomato+ adipocytes were seen in the Krt19^CreERT^ sample (**Fig. 3a**).

**Figure 3.**
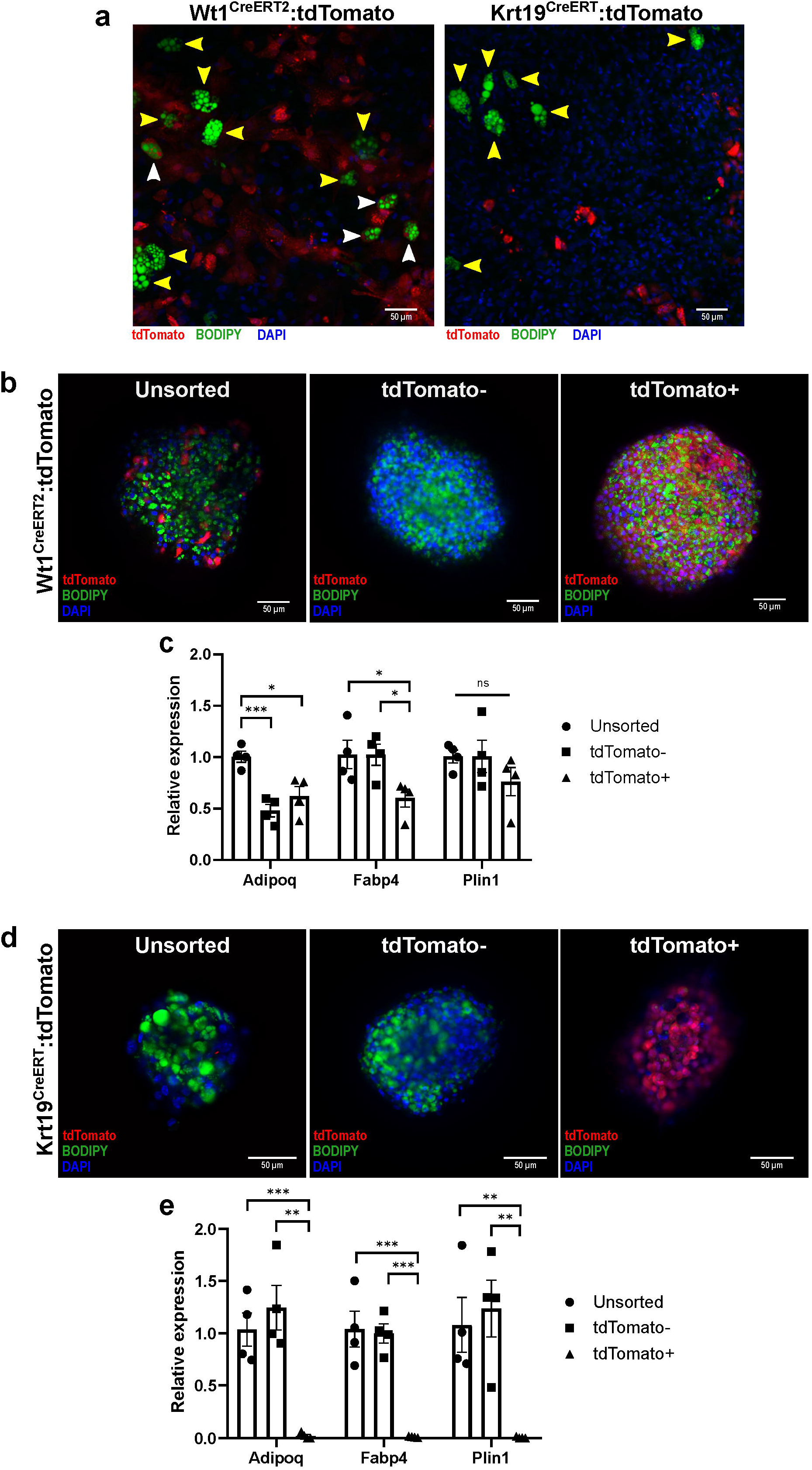
Krt19-expressing mesothelial cells do not differentiate into adipocytes *in vitro*. **a**, SVF from Wt1^CreERT2^ and Krt19^CreERT^ mice was isolated and underwent adipogenic differentiation on standard collagen-coated plates. While adipocytes differentiated from Wt1^CreERT2^:tdTomato SVF contained both tdTomato+ (white arrow heads) and tdTomato-(yellow arrow heads), no tdTomato+ adipocytes were found in the Wt1^CreERT2^:tdTomato samples. Representative images are shown. **b-c**, SVF was isolated from Wt1^CreERT2^:tdTomato mice after tamoxifen labeling, and was sorted into tdTomato+ and tdTomato-groups and plated on low-adhesion plates, forming microspheres. After adipogenic differentiation, unsorted, tdTomato+ and tdTomato-cells derived from Wt1^CreERT2^:tdTomato mice as demonstrated by BODIPY labeling of lipid droplet formation and qPCR for adipocyte markers *Adipoq, Fabp4*, and *Plin1*. **d-e**, SVF isolated from Krt19^CreERT^:tdTomato mice was sorted into tdTomato+ and tdTomato-populations, and while unsorted and tdTomato-cells differentiated normally as shown by BODIPY staining and qPCR, tdTomato+ cells did not undergo adipogenic differentiation. Error bars represent the standard error of the mean. ns, p ≥ 0.05, *, p <0.05; **, p<0.01; ***, p <0.001.

We next employed a 3D spheroid differentiation model in an effort to improve differentiation of visceral adipocytes from their progenitors (Emont et al., 2015), which also has the advantage of downscaling for use on only a few thousand cells (Spallanzani et al., 2019). Wt1^CreERT2^: and Krt19^CreERT^:tdTomato+ mice were injected with tamoxifen and VAT SVF was isolated a week later and sorted by flow cytometry into tdTomato+ and tdTomato-populations, which were subsequently plated on an ultra-low attachment surface plate to facilitate spheroid formation and then exposed to adipocyte differentiation media. While tdTomato+ cells isolated from Wt1^CreERT2^ mice differentiated into adipocytes, as demonstrated by BODIPY staining for lipid accumulation as well as qPCR for the adipocyte-related genes adiponectin (*Adipoq*), fatty acid-binding protein 4 (*Fabp4*), and perilipin 1 (*Plin1*) (**Fig. 3b-c**), tdTomato+ cells isolated from Krt19^CreERT^ mice did not accumulate lipid or express adipocyte genes (**Fig. 3d-e**).

### *Krt19*-expressing mesothelial cells do not contribute to mature adipocytes in visceral adipose depots *in vivo* in response to aging or high-fat diet

After demonstrating that *Krt19*-expressing mesothelial cells do not undergo adipocyte differentiation *in vitro*, we sought to confirm these findings *in vivo*. Wt1^CreERT2^: and Krt19^CreERT^ mice were crossed with the mTmG reporter model, in which Cre recombination results in a switch from tdTomato to GFP expression, and male and female offspring were labeled with tamoxifen at six weeks of age.

Littermates were placed on high fat diet one week after injection, or maintained on standard chow diet. At 15 weeks of age, mice were sacrificed and gonadal VAT was whole mounted for confocal microscopy (Berry et al., 2014). While there was GFP labeling of the mesothelial layer in both models (**Fig. 4a-b, d**), GFP+ adipocytes were only seen in Wt1^CreERT2^ fed either chow or HFD (**Fig. 4a-b**). Confocal images of 6-10 sections from 3 mice per group were obtained, and the percentage of GFP+ adipocytes was calculated (**Fig. 4c**). Despite counting over 2,200 adipocytes in the Krt19^CreERT^ chow-fed cohort and over 1,600 adipocytes in the HFD-fed Krt19^CreERT^ cohort, no GFP+ adipocytes were identified. Both male and female perigonadal depots were studied, with no difference noted between the sexes in either Wt1^CreERT2^ or Krt19^CreERT^ models (**Fig. S2a**). To further rule out the adipogenic potential of *Krt19*-lineage cells, we extended the period of high fat feeding to 18 weeks, but still saw no GFP+ adipocytes, and aging chow fed Krt19^CreERT^:mtmG for one year also failed to yield any GFP+ adipocytes (**Fig. S2b-c**).

**Figure 4.**
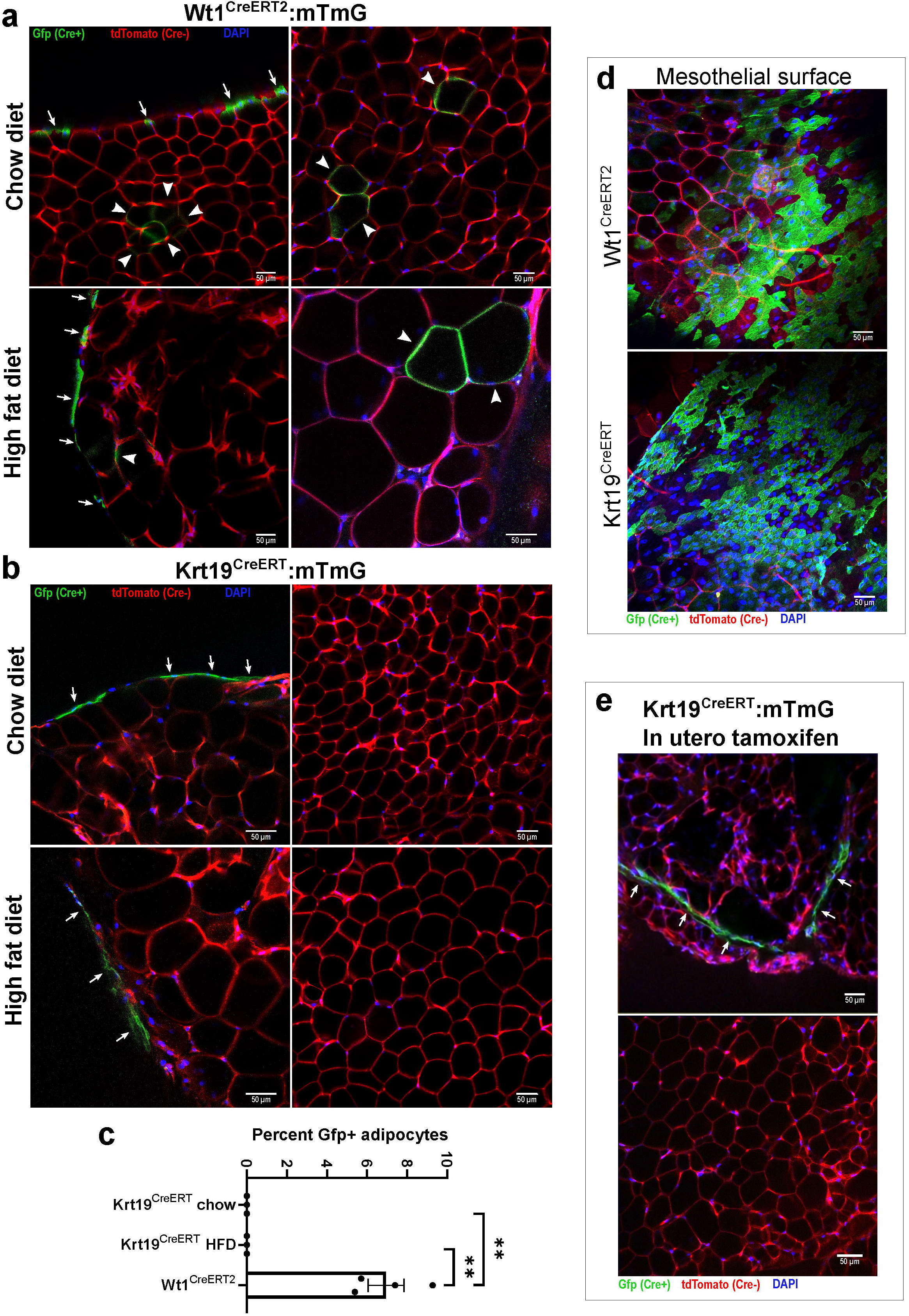
Krt19-lineage mesothelial cells do not differentiate into adipocytes in vivo. Wt1^CreERT2^:mTmG and Krt19^CreERT^:mTmG mice were injected with tamoxifen at 6 weeks of age and sacrificed at 15 weeks of age; mice receiving high fat diet did so starting at 7 weeks age. Whole mounts of eWAT were imaged by confocal microscopy. **a**, In Wt1^CreERT2^:mTmG mice, labeling of the mesothelial layer (arrows) as well as multiple Gfp+ adipocytes (arrow heads) were readily observed. **b**, In contrast, Krt19^CreERT^:mTmG mice had labeling only of the mesothelial layer and there were no GFP+ adipocytes observed. **c**, The percentage of Gfp+ adipocytes was quantified. In the chow and HFD Krt19-CreER groups, over 2,200 and 1,600 adipocytes, respectively, were counted without any Gfp+ adipocytes observed. **d**, GFP+ mesothelium was readily identifiable in both Wt1^CreERT2^:mTmG and Krt19^CreERT^:mTmG models. **e**, In order to evaluate the possibility that mesothelial cells which develop *in utero* may transition to a preadipocyte identity later in development, Krt19^CreERT^:mTmG dams were injected with tamoxifen at gestational day E14.5, and offspring sacrificed at 6 weeks of age. The mesothelium contained GFP+ cells, but there were no GFP+ adipocytes identified. Error bars represent the standard error of the mean. **, p<0.01.

Data derived from mice labeled with tamoxifen as adults do not preclude the possibility that the fetal mesothelium could give rise to adipose precursors which still express *Wt1*, even though the adult mesothelium may no longer contribute to the adipocyte pool. In fact, Chau, et al. demonstrated that exposing Wt1^CreERT2^:mTmG mice to tamoxifen in utero at day E14.5 resulted in GFP+ adipocytes in the adult mouse (Chau et al., 2014). We therefore repeated this experiment using Krt19^CreERT^:mTmG mice, exposing them to tamoxifen at day 14.5 of gestation via subcutaneous injection of the dam. This manipulation led to GFP+ cells in the mesothelium, but again we did not identify any mature GFP+ adipocytes in the grown offspring (**Fig. 4e**).

## Discussion

Adipose tissue development is a complex process that occurs in a depot-specific manner (Kahn et al., 2019). While some have suggested that white adipose tissue (WAT) adipocyte progenitors are homogenous and lack subtypes (Acosta et al., 2017), numerous studies now indicate that there is diversity within the preadipocyte pool (Ferrero et al., 2020). This complexity has led to challenges in identifying gene markers that comprehensively identify adipocyte progenitors, and almost a dozen different Cre lines have been used to perform embryonic adipocyte lineage tracing (Sebo and Rodeheffer, 2019). Beyond developmental preadipocyte commitment, terminal adipocyte differentiation from progenitors is also under the control of various metabolic and hormonal stimuli, including caloric overload (Wang et al., 2013), cold exposure (Wang et al., 2013), ischemia (Zangi et al., 2017), and wound healing (Plikus et al., 2017), adding to the complexity of studying adipocyte maturation. ScRNA-seq has provided an important tool to unravel the complexity of adipose tissue and generate hypotheses about its development and the role of different cell types under varying metabolic conditions (Ferrero et al., 2020).

The role of the mesothelium as a source of visceral preadipocytes during development and adulthood is intriguing and plausible given the variety of sources for adipocytes that have been previously identified. The idea has gained widespread acceptance, and subsequent studies have used *Wt1* as a marker for mesothelial-derived preadipocytes (Lee et al., 2019). Chau, et al. demonstrated that *Wt1*+ cells in visceral fat pads express the preadipocyte marker Sca-1, and argue that this is consistent with mesothelial cells’ capacity to undergo adipocyte differentiation (Chau et al., 2014). However, we demonstrate that Sca-1 and *Pdgfra* are in fact not expressed in the mesothelium, while *Wt1* is expressed in both the mesothelium and preadipocyte populations, indicating that *Wt1*+ mesothelial cells are distinct from *Wt1*+ preadipocytes. Using a tamoxifen-inducible Cre model driven by the specific mesothelial transcriptional marker *Krt19*, we demonstrate that mesothelial cells do not contribute to VAT adipogenesis in mice in homeostatic conditions or with caloric overload. That *Wt1* is expressed in visceral but not subcutaneous depots may indeed suggest distinct embryologic origins among WAT depots, but our findings clarify that the mesothelium does not represent a developmental intermediate for adipocytes.

The physiology of the mesothelium of visceral adipose depots remains under-characterized. Mesothelial cells have been suggested to play a role in adipose tissue inflammation in obesity (Darimont et al., 2008), and a recent study indicated the presence of an inflammatory subset of mesothelial cells that expands in mice after HFD feeding (Sárvári et al., 2021). *Wt1*-expressing preadipocytes and adipocytes have been previously described to be more highly-responsive to TNFα than other white adipocytes (Lee et al., 2019), and this could be taken to support the idea that inflammatory mesothelial cells may be differentiating into more pro-inflammatory adipocytes. However, given the distinction between *Wt1*+ mesothelium and *Wt1*+ preadipocyte pools demonstrated by our data, it is clear that studies of the mesothelium should use a more specific marker than *Wt1*, and studies of *Wt1*-expressing cells must recognize the underlying heterogeneity of that population.

In addition to elucidating key biology of the VAT mesothelium, this study raises important issues with respect to lineage tracing based on transcriptional markers. Identifying a single specific and sensitive marker for lineage tracing can be challenging. For example, the aP2-Cre was initially used as an adipocyte-specific-Cre, until it was found to be inefficient at labeling all adipocytes and also labeled some blood lineage cells and endothelial cells (Jeffery et al., 2014). Given *Wt1*’s broad expression during development, it is perhaps not surprising that it lacks specificity for a single cell type within a tissue, consistent with findings in other tissues in which *Wt1* is critical to development, such as the heart (Rudat Carsten and Kispert Andreas, 2012). Intersectional genetics approaches have been developed to address these challenges (Han et al., 2021), and the increasingly widespread application of single-cell and single-nucleus RNA sequencing will hopefully make the selection of cell-type specific markers more precise and more easily interpretable.

## Author contributions

Conceptualization, GPW, MPE, and EDR; Methodology, GPW, MPE, LT; Software, CJ and LT; Formal analysis, GPW, CJ and LT; Investigation, GPW, MPE, and JL. Writing – Original draft, GPW; Writing – Review & Editing, GPW, MPE, EDR, LT, CJ, and JL; Funding Acquisition, GPW, MPE, and EDR; Supervision, EDR; Project Administration, EDR.

## Acknowledgements

We gratefully acknowledge the Boston Nutrition Obesity Research Center Functional Genomics and Bioinformatics Core (NIH P30DK046200) for their expertise in single-cell RNA sequencing analysis, and the Beth Israel Deaconess Medical Center Flow Cytometry and Confocal Imaging Cores for their assistance in flow sorting and confocal microscopy studies, respectively. This work was supported by funding from the NIH (RC2DK116691). GPW received funding from NIH grant T32DK007516, administrative supplement RC2DK116691-03S1, and the Endocrine Fellows Foundation. MPE is supported by NIH grant F32DK124914. We thank all members of the Rosen lab for thoughtful and constructive discussion.

## STAR Methods

## RESOURCE AVAILABILITY

### Lead contact

Further information and requests for resources and reagents should be directed to the corresponding author Evan Rosen

### Materials availability

This study did not generate new unique reagents.

### Data and code availability

The datasets supporting the current study have not yet been deposited in a public repository but are available from the corresponding author upon request.

## EXPERIMENTAL MODEL DETAILS

### Animals

All animal experiments were performed in accordance with a protocol approved by the BIDMC Institutional Animal Care and Use Committee. Mice were maintained under a 12 hr light/12hr dark cycle at 22°C on chow diet (8664 Harlan Teklad, 6.4% wt/wt fat) unless otherwise specified. Krt19-CreERT(Means et al., 2008) (stock no. 026925), Wt1-CreERT2(Zhou et al., 2008) (stock no. 010912) lox-stop-lox tdTomato (stock no 007914) and mTmG mice (stock no 007676) were all obtained from Jackson Labs. Tamoxifen-induced Cre labeling was performed at 6 weeks of age with subcutaneous tamoxifen injections (100 mg/kg dissolved in sunflower seed oil) on four consecutive days. For high fat diet experiments mice were injected with tamoxifen at 6 weeks of age as above and either continued on chow diet or switched to high fat diet consisting of 20% calories from protein, 60% from fat, and 20% from carbohydrate (Research Diets, D12492) at 7 weeks of age; mice were sacrificed at 15 weeks of age for confocal imaging. For in utero tamoxifen exposure, the dam was injected with a single dose of tamoxifen (100 mg/kg) subcutaneously on day E14.5 of gestation; offspring were sacrificed at 6 weeks of age for confocal imaging. Both male and female mice were used to perform the presented experiments and similar results were obtained from mice of both sexes.

## METHOD DETAILS

### SVF isolation and flow cytometry

SVF isolation was performed as described elsewhere (Cho et al., 2014); briefly, mouse gonadal fat depots were washed with PBS on ice and then minced and incubated with collagenase type II (1 mg/ml) (Sigma-Aldrich) and incubated on a shaker water bath incubator at 37°C for 20 minutes with vigorous shaking by hand every 5 minutes. EDTA, pH 8.0 at a concentration of 10 mM was then added for an additional 5 minutes, after which wash media (DMEM+glutamax with 10% bovine calf serum and 1% penicillin-streptomycin) or FACS buffer (PBS with 1mM EDTA, 25 mM HEPES and 1% fetal bovine serum) was added. Cells were filtered through a 100 μm filter and washed, and if flow cytometry was to be performed, incubated with ACK lysis buffer for 5 minutes. SVF was then washed and filtered through 40 μm filter followed by plating on collagen-coated multiwell plates in growth media (DMEM+glutamax, 15% FBS, 1% penicillin-streptomycin) or suspended in FACS buffer for flow cytometry. For flow cytometry, cells were incubated in FITC-conjugated mouse anti-CD150a (PDGFRA) antibody (1:1000 dilution) (ThermoFisher 11-1401-82). Flow cytometric cell sorting and analysis was performed on a SORP FACSAria III (BD). Flow cytometry data was processed using FlowJo version 10.7.1 (BD).

### Trypsin mesothelial digestion

Gonadal fat pads were removed intact and washed once in PBS on ice and then transferred to trypsin solution (0.25% trypsin, 0.1% EDTA, Corning diluted 1:1 in PBS) and placed in shaker water bath incubator at 37°C for 30 minutes, with vigorous shaking by hand every several minutes. Intact adipose depots were then removed with forceps and minced and processed as described above (“remainder SVF”). Wash media was added to the remaining solution which contained mesothelial cells and cells were washed twice prior to plating. For qPCR studies, cells from individual animals were plated on separate wells of a collagen-coated 12-well plate for 1 hour, washed with PBS three times, and lysed in Trizol (Thermo Fisher). For confocal imaging, mesothelial cells from two mice were plated per well of a BioCoat collagen-coated 4-well culture slide (Corning) for approximately 16 hours prior to fixation with 4% paraformaldehyde. After fixation, cells were permeabilized with Triton X-100, stained with Hoechst, and coverslips mounted with Fluoromount-G (Southern Biotech).

### In vitro adipocyte differentiation assays

For standard in vitro differentiation assays, SVF was plated on collagen-coated 12-well plates in growth media for 2 days and then switched to differentiation induction media containing DMEM+glutamax, 10% FBS, 1% penicillin-streptomycin, 0.5 μg/ml insulin, 5 μM dexamethasone, 1 μM rosiglitazone, and 0.5 mM 3-isobutyl-1-methylxanthine for two days followed by maintenance media (DMEM+glutamax, 10% FBS, 1% penicillin-streptomycin, 0.5 μg/ml insulin) for an additional seven days prior to imaging, with media changes every 2 days. Cells were fixed with a formaldehyde-containing fixative Z-fix and stained with BODIPY to label neutral lipid accumulation that is consistent with adipocyte differentiation, and Hoechst for nuclear labeling. For microsphere differentiation, cells were flow-sorted as above, washed in wash media, and plated on collagen-coated multi-well plates for 16 hours. Cells were then suspended with trypsin and counted prior to plating approximately 5,000 cells per well into a 96-well ultra-low attachment surface plate (Corning, cat# 4515). Pre-plating the cells appeared to improve viability and spheroid formation compared to directly sorting into the ultra-low attachment surface plate. Cells were cultured in growth media for 2 days to allow spheroid formation, followed by 3 days in differentiation induction media as described above, followed by 5 days in maintenance media with insulin alone, with media change after 3 days in maintenance media. A subset of spheroids were then washed in PBS, fixed with Z-fix, stained with BODIPY and Hoechst, and placed onto slides for confocal imaging, while the remaining spheroids were washed and lysed in Trizol for RNA purification and qPCR.

### Quantitative PCR

Cells were collected in Trizol as above. Total RNA was isolated using RNeasy MinElute Cleanup Kit (Qiagen) using manufacturer’s instructions. Up to 1μg of RNA was reverse-transcribed using High-Capacity cDNA Reverse Transcription Kit (Thermo Fisher). SYBR Green PCR Master Mix (Thermo Fisher) and gene primers (**Supplementary table 1**) were used to perform qPCR on QuantStudio 6 Flex Real-Time PCR system (Thermo Fisher) and normalized to housekeeping gene Tbp. Results are presented as fold change (2^-ΔCT^).

### Confocal microscopy

All confocal images were obtained using a Zeiss LSM 880 Upright Laser Scanning Confocal Microscope using a 10X or 20X objective. Images were processed using Zen Black 2.3 software. Multi well slides and microspheres were prepared as above. Whole mount images were obtained by dividing gonadal depot into several small sections, washing in PBS once, fixing with Z-fix and Hoechst for 20 minutes, washing with PBS three times, and mounting on a slide with Fluoromount-G.

## QUANTIFICATION AND STATISTICAL ANALYSIS

### Single-cell RNA sequencing & clustering

Drop-seq was performed on Collagenase-digested adipose tissue SVF derived from 10-week old wild-type male C57BL/6 mice using procedures described in detail previously (Campbell et al., 2017; Macosko et al., 2015). Raw sequencing data were filtered for quality and trimmed of adapter sequences and poly(A) tails using Drop-seq tools (https://github.com/broadinstitute/Drop-seq), then aligned to the mouse genome (GRCm38/mm10) with STAR (Dobin et al., 2013) v2.6.1e. Digital expression matrices were generated from uniquely-mapped reads, including intronic and exonic features from the GENCODE M16 mouse genome annotation. Doublet cell populations were identified and removed with Scrublet (Wolock et al., 2019) using a doublet score threshold of 0.25. Clustering and further analyses were performed with the Seurat (v3.0.3.9036) software package (Satija et al., 2015; Stuart et al., 2019). First, we removed genes expressed in fewer than 2 cells, and cells with fewer than 400 UMIs or with greater than 10% mitochondrial content. Data were normalized and scaled and variable features detected using SCTransform (Hafemeister and Satija, 2019). Cells were then clustered using the top 50 PCs among the variable gene set. UMAP (Becht et al., 2018) projections were generated from these same PCs for visualization. Human adipose single-nucleus RNA-seq data were processed similarly to the above. Reads were mapped to the GRCh38/hg38 human genome assembly and counts generated from mappings to GENCODE 27 genome annotation (introns and exons).

### Statistical analysis

Statistics were performed on GraphPad Prism version 8. Paired two-tailed T tests were performed on mesothelium-remainder SVF qPCR samples, and unpaired two-tailed T tests were performed on microsphere qPCR samples and in vivo adipocyte counts.

**Table S1.**
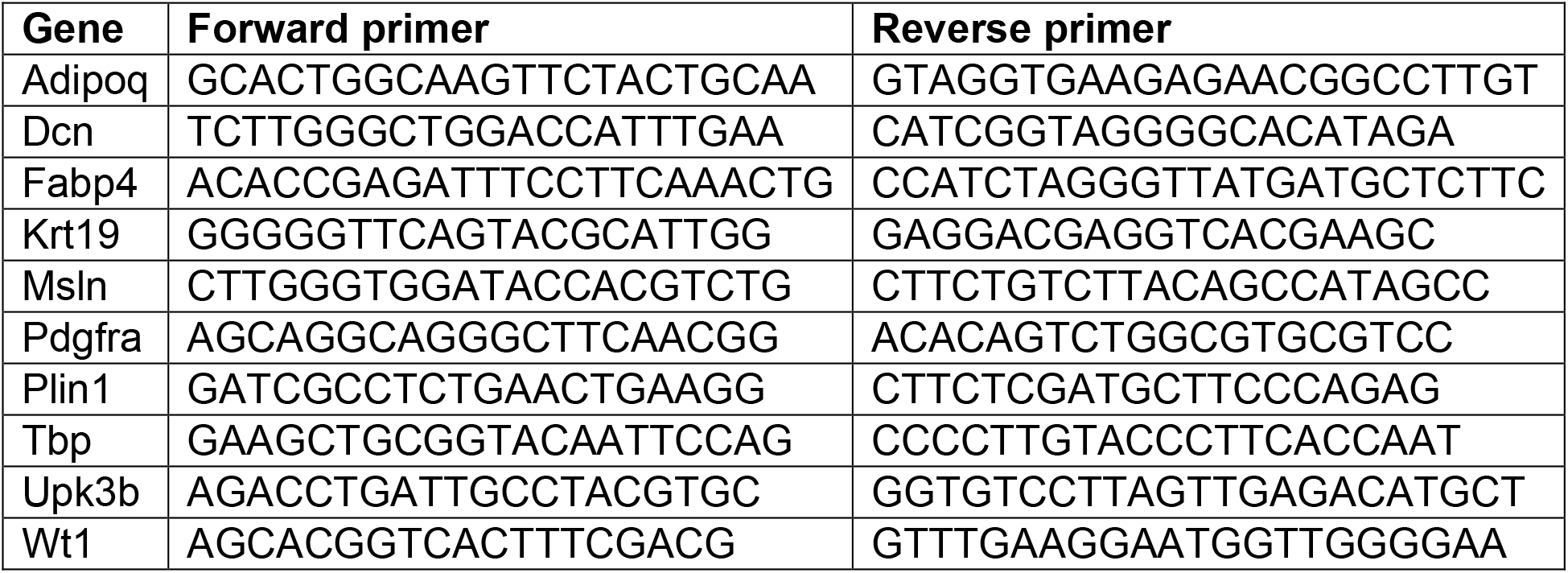
qPCR primer sequences

