## Supplemental figures for "Mesothelial cells are not a source of adipocytes in mice"

**Figure S1, related to Figure 1.** Wt1 is present in both preadipocyte and mesothelial populations in human omental adipose tissue

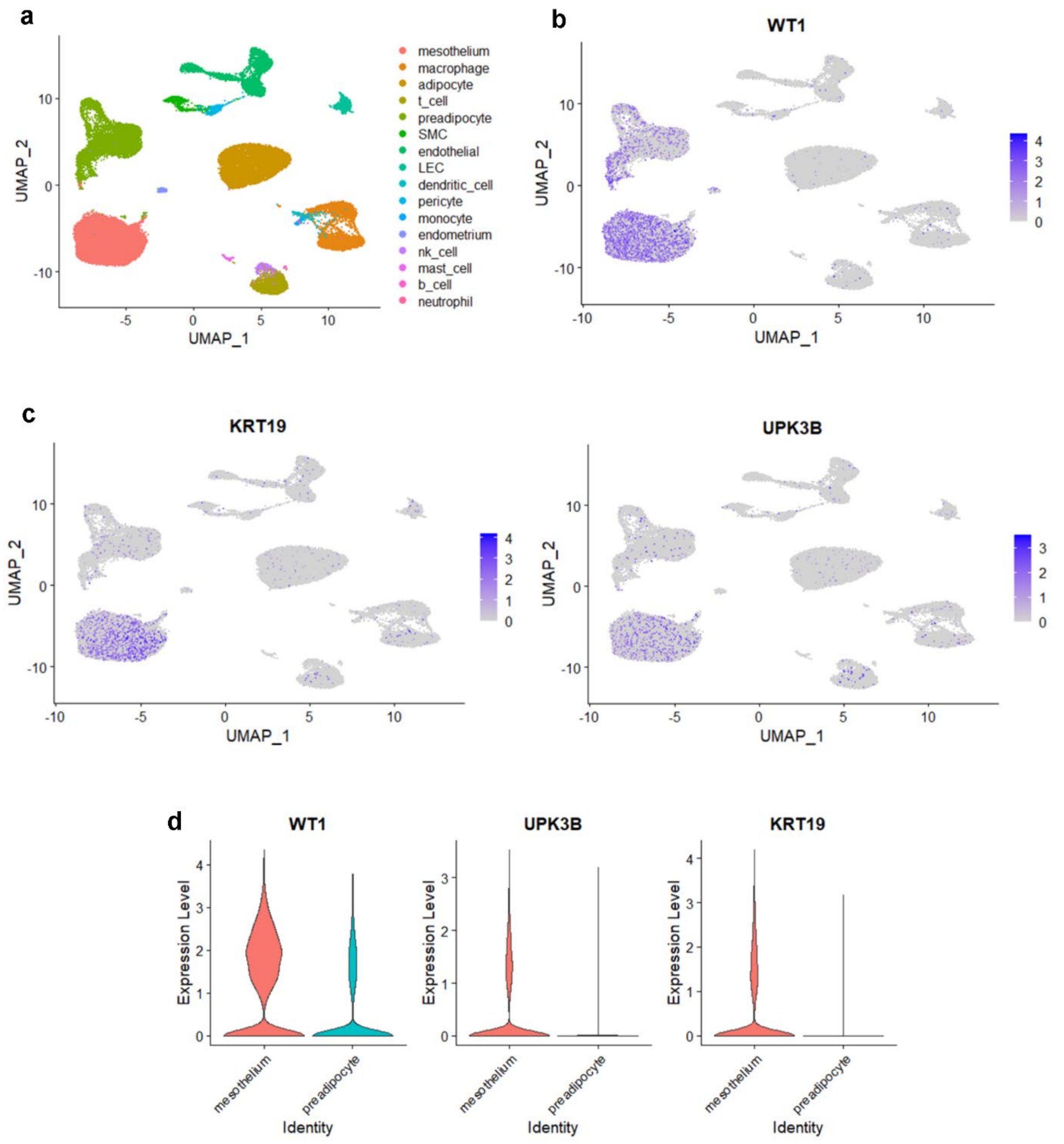


**a**, As in mouse, a UMAP of human omental single-nucleus RNA-sequencing data demonstrates distinct preadipocytes and mesothelial populations. **b**, WT1 is expressed in both preadipocyte and mesothelial populations. **c**, KRT19 and UPK3B are predominantly expressed in the mesothelial population only. **d**, Violin plots demonstrate WT1 expression in both mesothelial and preadipocyte populations, in contrast to specific mesothelial markers UPK3B and KRT19. SMC, smooth muscle cell; LEC, lymphatic endothelial cell.

**Figure S2, related to Figure 4.** Krt19+ mesothelial cells do not differentiate into adipocytes regardless of mouse sex, prolonged HFD, or aging

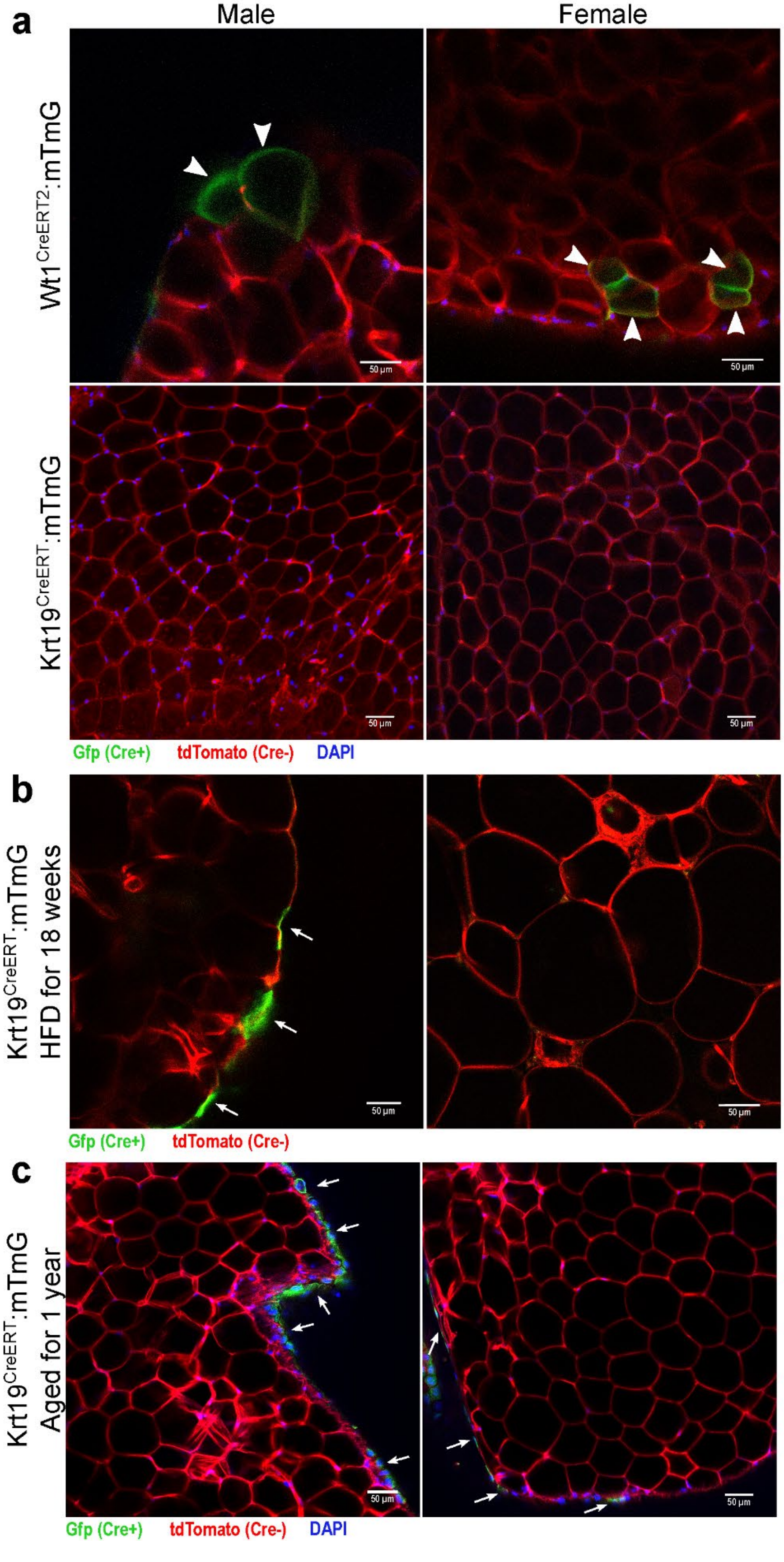


**a**, Both male and female perigonadal VAT depots were examined for the presence of GFP-labeled adipocytes. GFP+ adipocytes were identified in both male and female Wt1$^{CreERT2}$:mTmG VAT, while none were found in male or female

$Krt19^{CreERT}$:mTmG VAT. **b**, An 18-week course of HFD, and **c**, a year-long aging protocol failed to produce GFP+ adipocytes in labeled $Krt19^{CreERT}$:mTmG mice. Arrow heads: GFP-positive adipocytes; arrows: GFP labeling of the mesothelial layer. Representative images are shown.
